# Integrative Deep Models for Alternative Splicing

**DOI:** 10.1101/104869

**Authors:** Anupama Jha, Matthew R. Gazzara, Yoseph Barash

## Abstract

Advancements in sequencing technologies have highlighted the role of alternative splicing (AS) in increasing transcriptome complexity. This role of AS, combined with the relation of aberrant splicing to malignant states, motivated two streams of research, experimental and computational. The First involves a myriad of techniques such as RNA-Seq and CLIP-Seq to identify splicing regulators and their putative targets. The second involves probabilistic models, also known as splicing codes, which infer regulatory mechanisms and predict splicing outcome directly from genomic sequence. To date, these models have utilized only expression data. In this work we address two related challenges: Can we improve on previous models for AS outcome prediction and can we integrate additional sources of data to improve predictions for AS regulatory factors. We perform a detailed comparison of two previous modeling approaches, Bayesian and Deep Neural networks, dissecting the confounding effects of datasets and target functions. We then develop a new target function for AS prediction and show that it significantly improves model accuracy. Next, we develop a modeling framework to incorporate CLIP-Seq, knockdown and over-expression experiments, which are inherently noisy and suffer from missing values. Using several datasets involving key splice factors in mouse brain, muscle and heart we demonstrate both the prediction improvements and biological insights offered by our new models. Overall, the framework we propose offers a scalable integrative solution to improve splicing code modeling as vast amounts of relevant genomic data become available.

**Availability:** code and data will be available on Github following publication.

## 1 Introduction

A key contributor to transcriptome complexity is alternative splicing (AS): the joining together of different exonic segments of a pre-mRNA to yield different gene isoforms. The most common type of AS event in human and mouse is exon skipping where a fraction of the mRNA produced include an exon while others skip it. Thousands of such variations were found to be highly conserved and common between tissues. Overall, more than 90% of multi-exon human genes are alternatively spliced [10, 17] and splicing defects have been associated with numerous diseases. This has motivated detailed studies of AS variations across tissues, developmental stages, and malignant states [12]. These studies monitor mRNA expression at exonic resolution using RNA-Seq in a variety of experimental conditions, including knockdown (KD), knockout (KO), or over-expression (OE) of condition specific splicing factors (SF). Other experiments monitor binding affinity of splice factors using several similar protocols involving UV cross-linking of the factor to the RNA, followed by immunoprecipitation and sequencing of the bound RNA fragments (CLIP-Seq).

In parallel, the fact that splicing outcome is highly condition specific and regulated by many factors led to an effort to computationally derive predictive ‘splicing codes’: models that use putative regulatory features (e.g. sequence motifs, secondary structure) to predict splicing outcome in a condition specific manner (e.g. brain tissue) [2, 3]. Concentrating on cassette exons, the most common form of AS in mammals, these models aimed to predict percent splicing inclusion (PSI, Ψ) of the alternative exon, or changes of its inclusion (dPSI, ΔΨ). Such models have been used successfully to identify novel regulators of splicing events in disease associated genes, and predict the effect of genetic variations on splicing outcome [7, 18, 14]. However, given the sharp growth in sequencing data, two main questions are: Can we leverage the new CLIP-Seq and splice factors KD/OE experiments and more generally, can we improve on current splicing code models?

Previous work has shown that Bayesian Neural Networks compare favorably to a plethora of other modeling approaches including KNN, SVM, Naive Bayes, and Deep Neural Networks with Dropouts [19, 15]. specifically, [15] described dropout as performing an approximation to the BNN Bayesian model averaging, and pointed to the latter as being advantageous for smaller datasets. However, later work using a Deep Neural Network with an autoencoder demonstrated improved performance compared to a BNN model [9]. Notably, these different works used different datasets and mixed the effect of modeling framework (BNN vs DNN) with changes of the target function. Thus, in this work we First reconstructed previous BNN and DNN models on the original dataset from [9]. After establishing these as a baseline, we then monitored the effect of a new target function, of increasing dataset size by exploiting improvements in RNA-Seq quantification algorithms [16], and adding new types of experimental data.

The First contribution of this work is in developing a new target function for splicing code models. Due to limitations of both available data and algorithms, previous works were unable to predict Ψ or ΔΨ directly. Instead, they formulated a three way prediction task 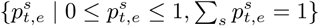 for any exon *e* in each condition *t*. In the original formulation, *s* represented the chances for increased inclusion, exclusion, or no change for exon *e* in condition *t*, compared to a hidden baseline of inclusion inferred from a set of 27 tissues [3]. This formulation allowed the learned model to concentrate its predictive power on tissue regulated exons, using a dedicated sparse factor analysis model to identify those exons from noisy micro-array data [2]. Subsequently, the same target function formulation was used, but instead of inferring splicing changes, *s* now represented binning of Ψ values into three levels: “Low” (0 ≤ Ψ < 33%), “Medium” (33% ≤ Ψ < 66%), and “High” (66% ≤ Ψ ≤ 100%). While useful, these target functions are inherently unsatisfying as an approximation to the underlying biological variability. Here, we develop a new target function which models Ψ directly, and demonstrate its improved accuracy compared to previous approaches. Serving as a baseline, Figure 1 depicts the improvement in percent variance explained by the new model compared to previous ones on the original dataset used by [9].

**Figure 1:**
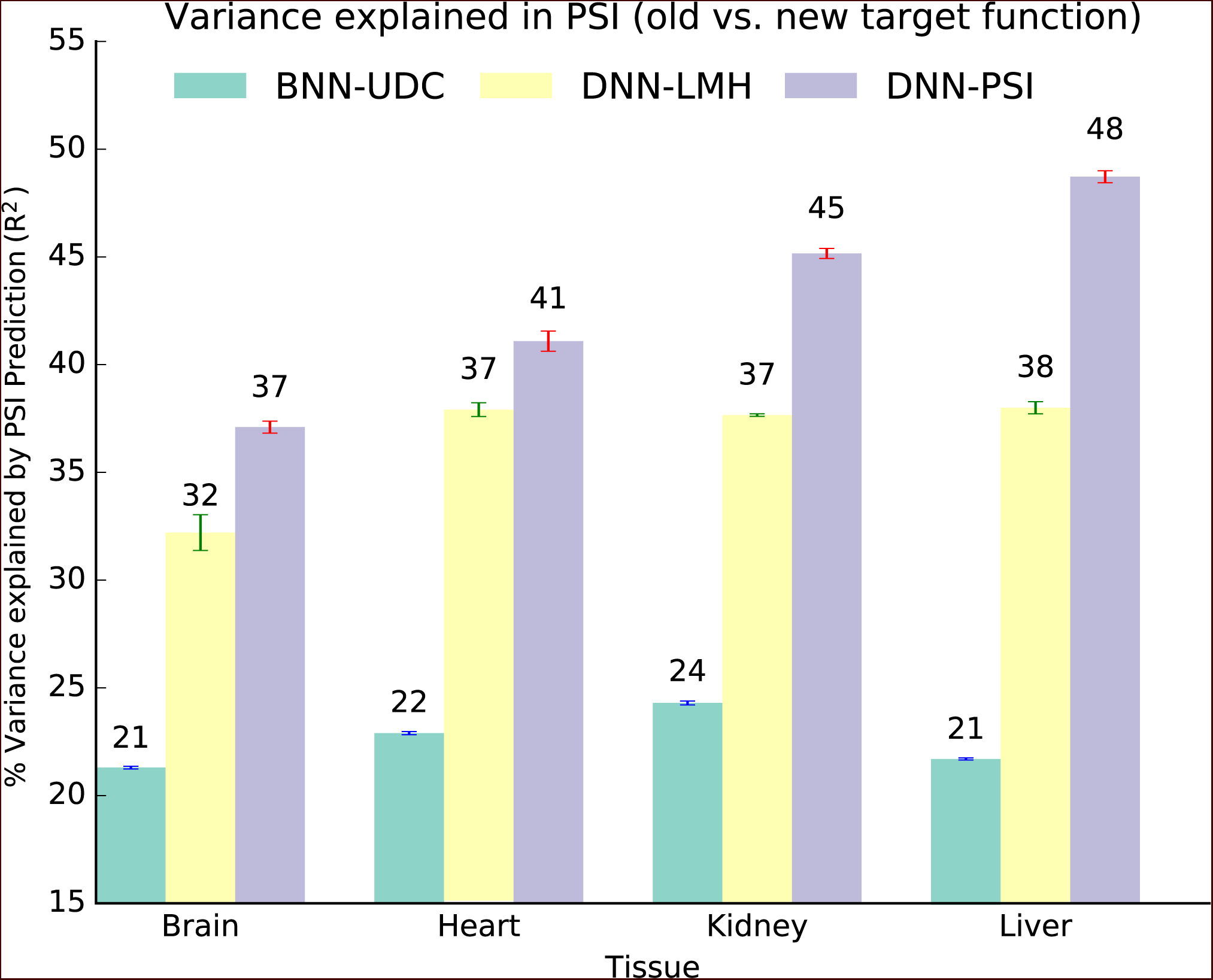
Improvement in %variance explained by the new target function (light purple) compared to previous BNN and DNN models on the original tissue data used by [9].

The second contribution of this work is developing a framework to integrate additional types of experimental data into the splicing code models. Specifically, CLIP-Seq based measurements of *in vivo* splice factors binding are turned into an additional set of input features while knockdown and over-expression experiments are added with binary vectors coding which tissue and which factor (if any) are measured. A graphical representation of the various old and new model architectures is given in Figure 2. We demonstrate the effect of the new integrative modeling approach using a set of CLIP-Seq, knockdown and overexpression experiments for members of the Rbfox, Celf, and Mbnl splice factors in mouse heart, muscle and brain. Finally, we showcase some of the possible biological usage cases for these splicing code models for accurate *in silico* prediction of splice factors KO effect, and for identifying novel regulatory interplay between different splice factors.

**Figure 2:**
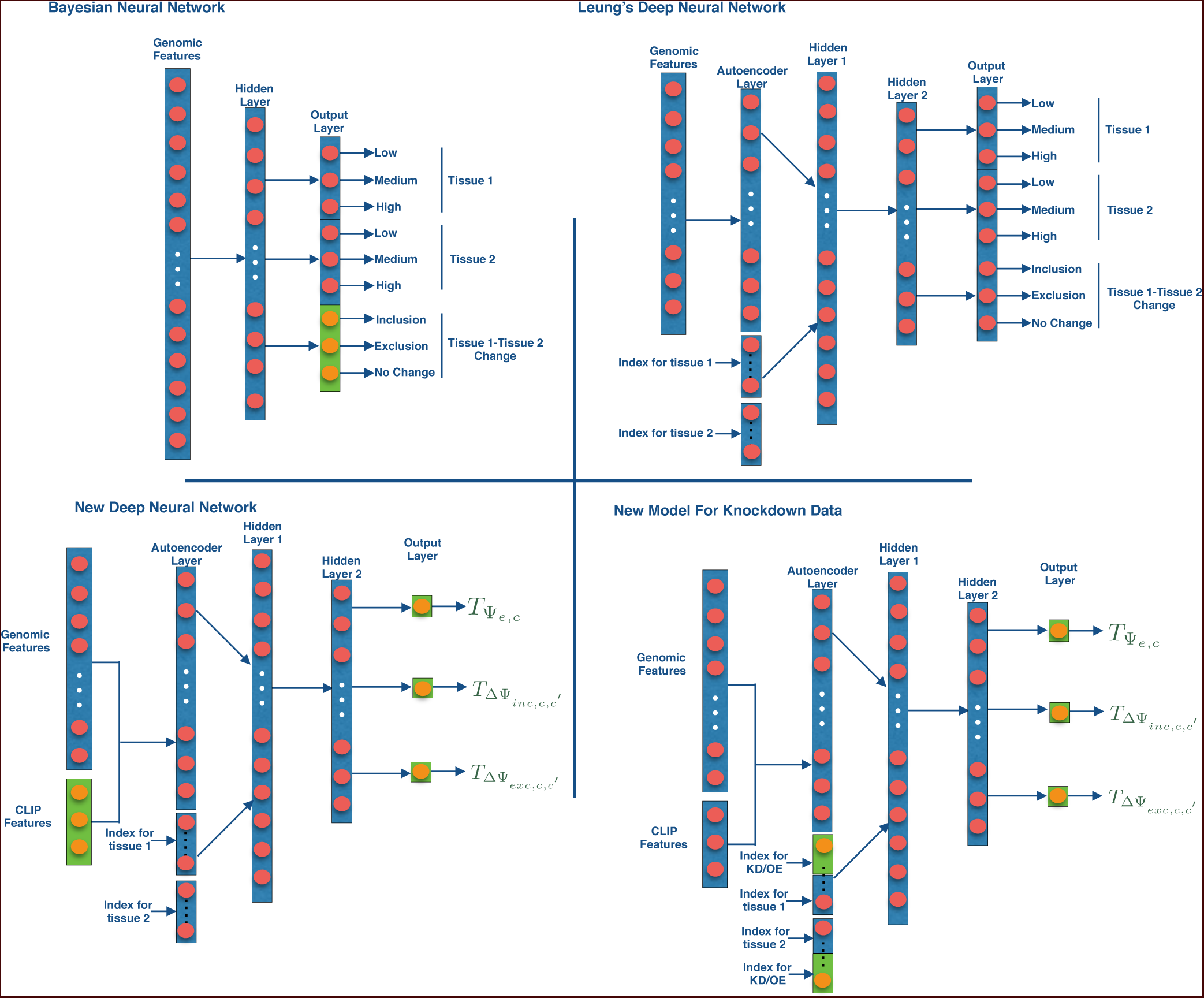
Architecture of the Bayesian Neural Network, old and new Deep Neural Network models

## 2 Methods

### 2.1 Datasets

Two RNA-Seq datasets were processed for this work. One, denoted Five Tissue Data, is the RNA-Seq data from five mouse tissues (brain, heart, kidney, liver and testis) produced by [5]. This dataset was used in the [9] paper and thus we use it to compare the old and new models. We generated genomic features and PSI quantification for approximately 12,000 cassette exons used in [4] for this dataset for the five tissues using MAJIQ [16] and AVISPA [4]. The second dataset, denoted MGP Data, was prepared by [8] and it contains RNA-Seq data from six tissues (heart, hippocampus, liver, lung, spleen, thymus) with average read coverage of 60 million reads. To this data we added 15 CLIP-Seq experiments (See Supplementary Table 10). Together, these datasets highlight some of the challenges involved in utilizing such diverse experiments. First, CLIP-Seq experiments give noisy measurement of where a splice factor binds. The measurements are noisy since binding signal (reads aligning to a certain area) may be false positives, may not indicate active regulation and may suffer from false negatives due to low coverage, indirect binding, antibody sensitivity, etc. Moreover, these experiments are typically executed by different labs, in different conditions and at varying levels of coverage. Thus, it is crucial that any learning framework that we develop should be able to handle missing and noisy measurements.

In our learning setting, the CLIP-Seq data is turned to input features indicating possible binding in a region proximal to the alternative exon (e.g. upstream intron). The target in our problem formulation is the relative exon inclusion level in a given experiment, expressed as percent spliced in (PSI, Ψ ∈ [0,1]). PSI serves to capture the proportion of isoforms that include the exon versus those that skip it. But since these are not observed directly, the short sequencing reads are used to construct a posterior beta distribution over PSI per exon 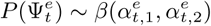. Similarly, when comparing two conditions the short reads are used to construct a posterior distribution over dPSI ΔΨ ∈ [-1,1] [16]. In practice many exons tend to be either highly included or highly excluded in any given condition, but approximately 20% of the measurements in our dataset have 0:1 < *E*[Ψ] < 0.9 and the concentration of the posterior Ψ or ΔΨ distribution around that mean value depends on the total number of reads hitting that region and how these are distributed across the transcriptome[16].

To enable comparison to previous works, we derived a feature set that excluded the additional CLIP based features described above. The 1,357 non CLIP-Seq features comprised of binary, integer and real valued features. These features have vastly different distributions with some being highly sparse, and some features being highly correlated (e.g. alternative representations of a splice factor binding motif). Finally, in any given condition only a small subset of those features are expected to represent relevant regulatory features.

Since many splicing changes occur in complex/non-binary splicing events, limiting the splicing code model to the original predefined 12,000 cassette events means that we may lose many important splicing variations. To capture additional cassette or cassette-like splicing variations we developed a pipeline that parses gene splice graphs constructed by MAJIQ to find additional training samples in the dataset. This process allowed us to find 2; 876 more events changing in at least one tissue comparison in the MGP data.

Next we processed seven splice factor knockdown, knockout and over-expression RNA-Seq datasets for four key splicing factors Celf1/2, Mbnl1 and Rbfox2 (for details of the datasets, see Supplementary Table 11). These datasets pose a challenge for any integrative learning framework since they are low coverage, noisy and generated by different labs.

We divided our datasets into 5 folds. Three folds were used for training, one for validation and one for testing. We repeated the modeling tasks 3 times, permuting the dataset each time to produce standard deviation estimates in the performance evaluation.

### 2.2 Likelihood Target Function

Motivated by the high noise in microarrays and later applied to RNA-Seq data, previous works translated the measurements of exon inclusion levels into a posterior distribution over random variable 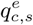 for each exon *e* and condition *c* with three possible assignments 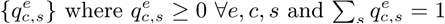. For PSI prediction, *s* ∈{*L,M,H*} represent chances of 0 Ψ ∈ < 0.33,0.33 ≤ Ψ < 0.66 and 0.66 ≤ Ψ ≤ 1 respectively. For changes in PSI, 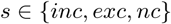 represent chances of increased inclusion, exclusion or no change. Consequently, an information theoretic code quality measure 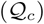 was used to score the predictions made by the splicing code. 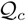
 is expressed as the difference in the Kullback-Leibler (KL) divergence between each target and predicted distribution:

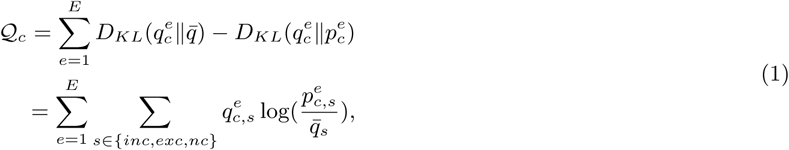

where *c* is the splicing condition (*e.g*., CNS), *E* is the number of exons and 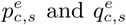 are the predicted and target probabilities. Alternatively, 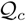 can be interpreted as the log-likelihood of the predictions minus the log-likelihood of a naive predictor based on the marginal distribution only.

Although useful, this target function suffers from several deficiencies when applied to RNA-Seq data. First, the binning process results in a rudimentary estimation of Ψ and ΔΨ. Second, the optimization only aims to bring 
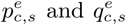
 closer, without any relation to order or meaning. For example, if a cassette event has low inclusion (*q_c,s=L_* ∼ 1) then predicting 
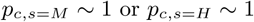
are just as bad. Moreover, in cases where an event suffers from insufficient or highly variable read coverage we may have 
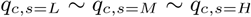
. In such cases, a model with high confidence (e.g. *p_c,s_=H* ∼ 1) based on sequence features will be penalized just the same, though there was no substantial evidence against it.

In order to overcome the above limitations, for every pair of conditions *c* and *ć*, we define three target variables:

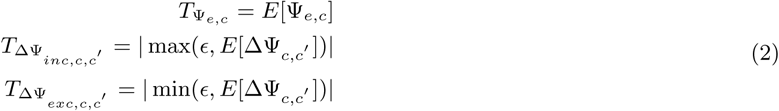

where 
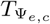
 is the expected PSI value of the event e in condition *c*. 
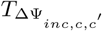
 captures the dPSI for events with increased inclusion between condition *c* and *ć* and 
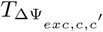
 captures the dPSI for events with increased exclusion between condition *c* and *ć*. ∈ is a uniform random variable with values between 0.01 and 0.03, it is used to provide very low dPSI values for non-changing events. 
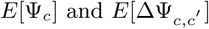
 were computed from the raw RNA-Seq data from condition *c* and *ć* using MAJIQ [16]. Given the above target variables definition, we define the new likelihood target function as

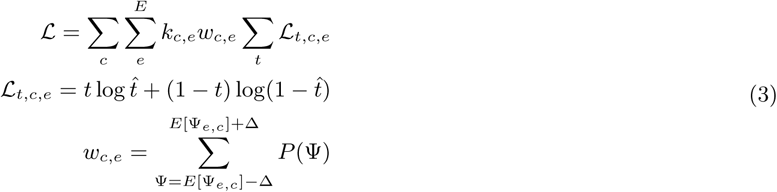

where 
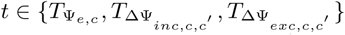
 and *k*_*c,e*_ = 1 if exon *e* is quantifiable in condition *c*. The weight *w*_*c,e*_ is defined by the probability mass in the area ±Δ around the expected ψ_*c*_ as defined by MAJIQ. This definition carries several benefits. First, it allows us to combine many different datasets, where the same event may or may not be quantifiable. Second, even when an event is deemed quantifiable (*k*_*c,e*_ = 1), the model can take into account how sure MAJIQ is in the ψ inferred from the RNA-Seq experiment.

### 2.3 Models

#### 2.3.1 Architecture

The BNN model was described in detail in [19, 9]. Briefly, the network consists of one hidden layer with 12 hidden units and sigmoidal non-linearity was used for each hidden unit. Network weights are random variables with a Gaussian distribution and a spike and slab prior which encourages sparsity. Figure 2 shows the network architecture of BNN used in this work. We note that [9] only used the LMH variables for the BNN. We supplemented those with UDC variables to make the BNN targets equivalent to those of the DNN architecture used in that work, leading to improved performance for the BNN model (see Supplementary Table 4).

The original DNN model shown in Figure 2 included an autoencoder layer with tanh activation and two hidden layers with ReLU activation units. Additionally tissue type was input as two, 5 (number of tissues) hot vectors where each bit represents a tissue and is active when the network is input an event comparing that tissue with another. For example, if the tissue order is [brain, heart, kidney, liver, testis] and the current comparison is brain versus heart, then the two tissue type hot vectors will be [1 0 0 0 0] and [0 1 0 0 0]. Dropout with probability 0.5 was used in each layer except the autoencoder layer. The hyperparameters are described in Supplementary Table 12. We experimented with different types of network architectures with different number of hidden layers and hidden units, different activation units and batch normalization. Since non of those architectures performed significantly better (data not shown) we decided to maintain the original DDN architecture for the purpose of this work.

The new DNN model architecture shown in Figure 2 includes the following additions. First, the target function has been changed as described in Section 2. We also added 874 CLIP features to the input dataset. We maintained the three layer structure of the original DNN models since we observed that adding additional layers did not improve performance. Batch normalization was performed at the second and third hidden layer and dropout with probability 0.5 was applied to both. We noticed that adding L1/L2-regularization did not have any impact on the model performance and we decided to exclude it from the final model. We allowed the learning rates of the three target variables to vary to capture optimal model performance.

As shown in Figure 2, for splice factor modeling, we modified the tissue type input to include the splice factor knockdown/knockout or over-expression. We used two 4 (number of tissues) hot vectors to represent the tissues and two 4 (number of splice factors) hot vectors to represent the splice factors. Since the datasets for this model were lower coverage and more noisy than the previous models, this model was more sensitive to different hyperparameter values during the tuning phase with cross validation. Three hidden layers were found to be optimal and L1-regularization was performed on the autoencoder layer. Dropout of 0.5 was used for the second and third hidden layers.

#### 2.3.2 Learning

Following the procedure suggested by [9], we trained the first layer of the model as an autoencoder for dimensionality reduction. This procedure proved beneficial for the new models as well. Next, the set of weights from the first layer were fixed and the tissue input was added. In the second stage, the two layered feed forward neural network was trained using SGD with momentum and weights were fine tuned by backpropagation. Each sample input to the network consists of 1,357 genomic (+ 874 CLIP) features and has three target variables, 
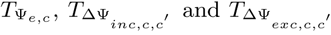
. Training batches are biased to prioritize changing events. Early stopping and dropout layers prevent the network from overfitting.

The three target variables are different in nature since one learns the baseline PSI and the other two learn the inclusion and exclusion dPSI. Thus, varying the learning rates for them optimizes the model performance on each one. The autoencoder network was trained for 300-500 epochs and the feed-forward neural network was trained for 1,000-1,400 epochs. Validation data was used for the hyperparameter tuning and once the set of hyper parameters were fixed, the final model was trained with the training and the validation data. 15 models were trained with the 5 folds and 3 permutations of the whole datasets. The performance evaluation is on the concatenated predictions of the test set from the 5 folds and error bars are computed using the 3 permutations. Tensorflow was used to develop the deep model and GPUs were used to accelerate the training process.

For the BNN, each tissue pair was trained as an independent model. Spike and slab prior was used to enforce sparsity and the weights were assumed to have a Gaussian distribution. 950 samples from the posterior distribution of weights were generated using 1,000 MCMC training iterations with Gibbs sampling. Initial 50 samples were discarded as burn-in. The final predictions are generated by averaging over the predictions from the 950 sampled weights. 15 models were trained per tissue comparison with 5 fold cross validation and 3 data permutations. After fixing the model hyperparameters, the validation data was included in training the final model.

## Results

For assessing the prediction accuracy, two types of measures have been used in this work. The predicted 
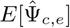
is compared to the estimated 
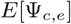
from the RNA-Seq experiments to compute the fraction of variance explained (*R*^2^). Area under the ROC curve (AUC) was computed for the predictions of exons that were differentially excluded/included 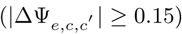 or not changing 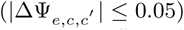.

We aim to measure the effect of each new element on the prediction accuracy. As a baseline, Figure 1 shows the effect of new target function on prediction with no other modeling additions on the original dataset used by [9]. We see significant improvement ( 10% - 25%) in PSI estimation and splicing target prediction (See Supplementary Table 6) by the new model (DNN-PSI) when compared to the DNN (DNN-LMH) and BNN (BNN-UDC) with the old target function. We added inclusion, exclusion and no change output variables for the Bayesian Neural Network since it improved splicing target prediction performance compared to the BNN without these labels (BNN-MLR [9],see Supplementary Table 4). DNN-LMH was designed according to the architecture and hyperparameters described in [9]. We note that the results for the previous models are not directly extracted from [9] but rather reconstructed to produce similar performance since both code and data were not available from the original publication. Also, since the DNN-LMH does not predict PSI directly, we computed the *E*[Ψ] as the weighted average of the {*L, M, H*} class prediction probabilities, following [18].

As noted earlier, previous works [4, 9, 18] were performed on a predefined set of approximately 12,000 alternative cassette exons. This approach of using only predefined cassette exons can limit the performance of the learned models, especially those involving deep neural networks which require large datasets. Thus, we developed a process termed cassettization (see Section 2.1) to detect and quantify additional cassette and cassette like alternative exons from RNA-Seq data. Also, due to the limited coverage of [5], we performed subsequent analysis on the MGP six tissues data described in Section 2.1. To assess the effect of cassettization on performance, we used two identically configured BNN models and trained one on the original 12,000 cassette exons (BNN-UDC) while the second got an additional training set with the cassettized events. Figure 3(a) shows that cassettization caused a substantial improvement in PSI estimation and splicing target prediction (See Supplementary Table 7) with all other factors being constant.

**Figure 3:**
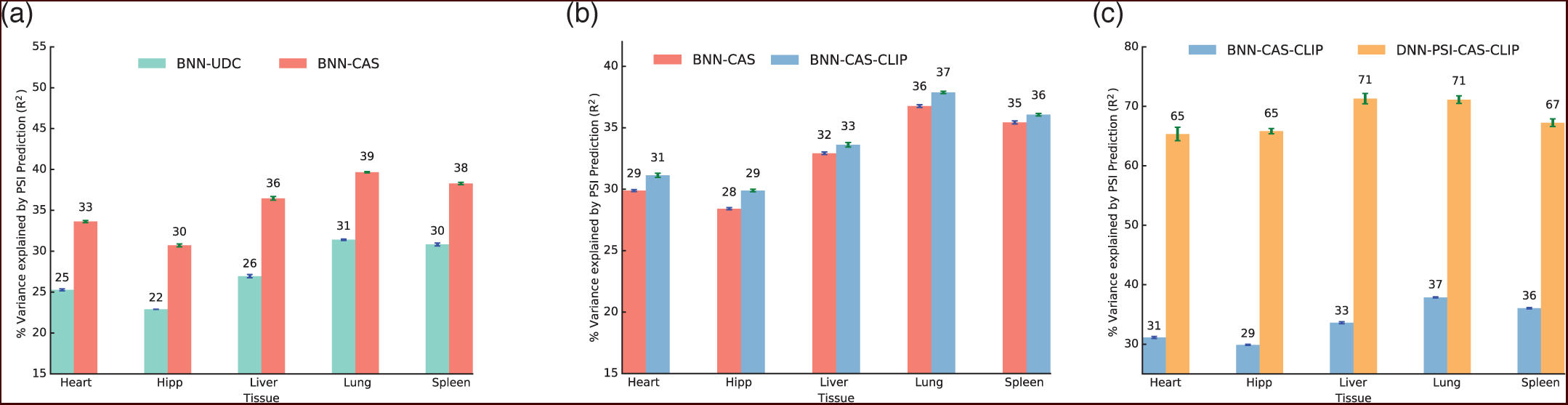
(a) effect of cassettization on PSI estimation. (b) effect of CLIP data on PSI estimation. (c) Comparison of old and new models with cassettization and CLIP

Our next goal was to measure the effect of CLIP-Seq data on PSI estimation. Using the same setup described above, we trained two BNNs identical in every aspect except one was given the CLIP data as input features (BNN-CAS-CLIP) and the other was not (BNN-CAS). Introducing CLIP features added a modest improvement to the PSI estimation as seen in Figure 3(b). One possible explanation for the modest improvement could be underfitting of BNN-CAS-CLIP since CLIP is introduced as new features to the model but the model’s hidden layer size and other hyperparameters are fixed.

In order to test the combined effect of the new target function, CLIP data and cassettization on the model’s performance and to compare BNN and DNN frameworks for the task of PSI estimation, we trained a BNN model with the old target function, cassettization and CLIP (BNN-CAS-CLIP) and a DNN model with the new target function, cassettization and CLIP (DNN-PSI-CAS-CLIP). Figure 3(c) and Table 1 summarize the results for the two model for both PSI estimation and splicing target prediction. Figure 3(c) shows large performance improvement of the new model for PSI estimation when compared to the BNN. This improvement carries over to the task of splicing target prediction seen in Table 1 as well, and for every tissue pair.

**Table 1:** Comparison of splicing target prediction of DNN-PSI-CAS-CLIP vs. BNN-CAS-CLIP in terms of AUCs of inclusion vs. all, exclusion vs. all and change vs. no change.

| Tissue Pair | Model | Inclusion | Exclusion | No Change |
| --- | --- | --- | --- | --- |
| Heart-Hipp | BNN-CAS-CLIP | 92.97±0.12 | 88.22±0.16 | 92.26±0.06 |
|  | DNN-PSI-CAS-CLIP | **95.70±0.06** | **94.09±0.34** | **94.72±0.06** |
| Heart-Liver | BNN-CAS-CLIP | 78.09±0.49 | 89.38±0.24 | 85.13±0.15 |
|  | DNN-PSI-CAS-CLIP | **92.15±0.60** | **96.26±0.18** | **94.11±0.26** |
| Heart-Lung | BNN-CAS-CLIP | 82.52±0.67 | 89.77±0.18 | 87.94±0.18 |
|  | DNN-PSI-CAS-CLIP | **92.15±0.80** | **95.42±0.30** | **93.60±0.26** |
| Heart-Spleen | BNN-CAS-CLIP | 79.37±0.21 | 91.03±0.13 | 87.45±0.08 |
|  | DNN-PSI-CAS-CLIP | **93.18±0.22** | **96.98±0.47** | **95.22±0.33** |
| Heart-Thymus | BNN-CAS-CLIP | 82.01±0.64 | 86.20±0.24 | 85.91±0.23 |
|  | DNN-PSI-CAS-CLIP | **92.76±0.36** | **95.83±0.15** | **94.06±0.32** |
| Hipp-Liver | BNN-CAS-CLIP | 83.33±0.08 | 93.16±0.02 | 90.32±0.07 |
|  | DNN-PSI-CAS-CLIP | **94.36±0.41** | **97.33±0.24** | **95.60±0.07** |
| Hipp-Lung | BNN-CAS-CLIP | 84.19±0.23 | 92.71±0.05 | 90.61±0.04 |
|  | DNN-PSI-CAS-CLIP | **93.32±0.33** | **95.92±0.11** | **94.47±0.16** |
| Hipp-Spleen | BNN-CAS-CLIP | 83.84±0.34 | 93.36±0.06 | 90.75±0.10 |
|  | DNN-PSI-CAS-CLIP | **93.77±0.09** | **96.86±0.13** | **95.51±0.10** |
| Hipp-Thymus | BNN-CAS-CLIP | 83.10±0.36 | 88.63±0.15 | 87.83±0.18 |
|  | DNN-PSI-CAS-CLIP | **91.77±0.27** | **95.64±0.10** | **94.46±0.05** |
| Liver-Lung | BNN-CAS-CLIP | 84.60±0.36 | 81.73±0.37 | 83.07±0.42 |
|  | DNN-PSI-CAS-CLIP | **98.14±0.54** | **94.23±0.15** | **95.71±0.28** |
| Liver-Spleen | BNN-CAS-CLIP | 85.41±0.40 | 87.66±0.15 | 87.59±0.21 |
|  | DNN-PSI-CAS-CLIP | **97.01±0.61** | **94.46±0.29** | **96.04±0.63** |
| Liver-Thymus | BNN-CAS-CLIP | 84.25±1.10 | 74.23±0.03 | 77.03±0.14 |
|  | DNN-PSI-CAS-CLIP | **96.80±0.76** | **93.27±0.38** | **93.44±0.20** |
| Lung-Spleen | BNN-CAS-CLIP | 79.82±0.32 | 80.71±0.49 | 80.71±0.09 |
|  | DNN-PSI-CAS-CLIP | **96.83±0.75** | **96.03±1.13** | **96.91±0.39** |
| Lung-Thymus | BNN-CAS-CLIP | 79.97±0.41 | 78.41±0.44 | 79.57±0.30 |
|  | DNN-PSI-CAS-CLIP | **94.98±1.05** | **96.51±0.47** | **95.91±0.16** |
| Spleen-Thymus | BNN-CAS-CLIP | 70.55±1.31 | 70.73±1.14 | 70.99±0.76 |
|  | DNN-PSI-CAS-CLIP | **97.91±0.47** | **91.86±1.23** | **92.21±0.85** |

Next, we turned to the new integrative framework that incorporates knockdown/knockout and over-expression experiments (see Section 2.3.1, Section 2.1). Figure 4(a) shows that the new integrative deep model generalizes well for this new type of KD/KO/OE data, offering large performance improvement for PSI estimation. One exception is the model performance on Rbfox2 KD in C2C12 cells. This may be due to the different experimental condition (C2C12 cells) or the number of samples, which require specific adjustments of the model’s training parameters.

**Figure 4:**
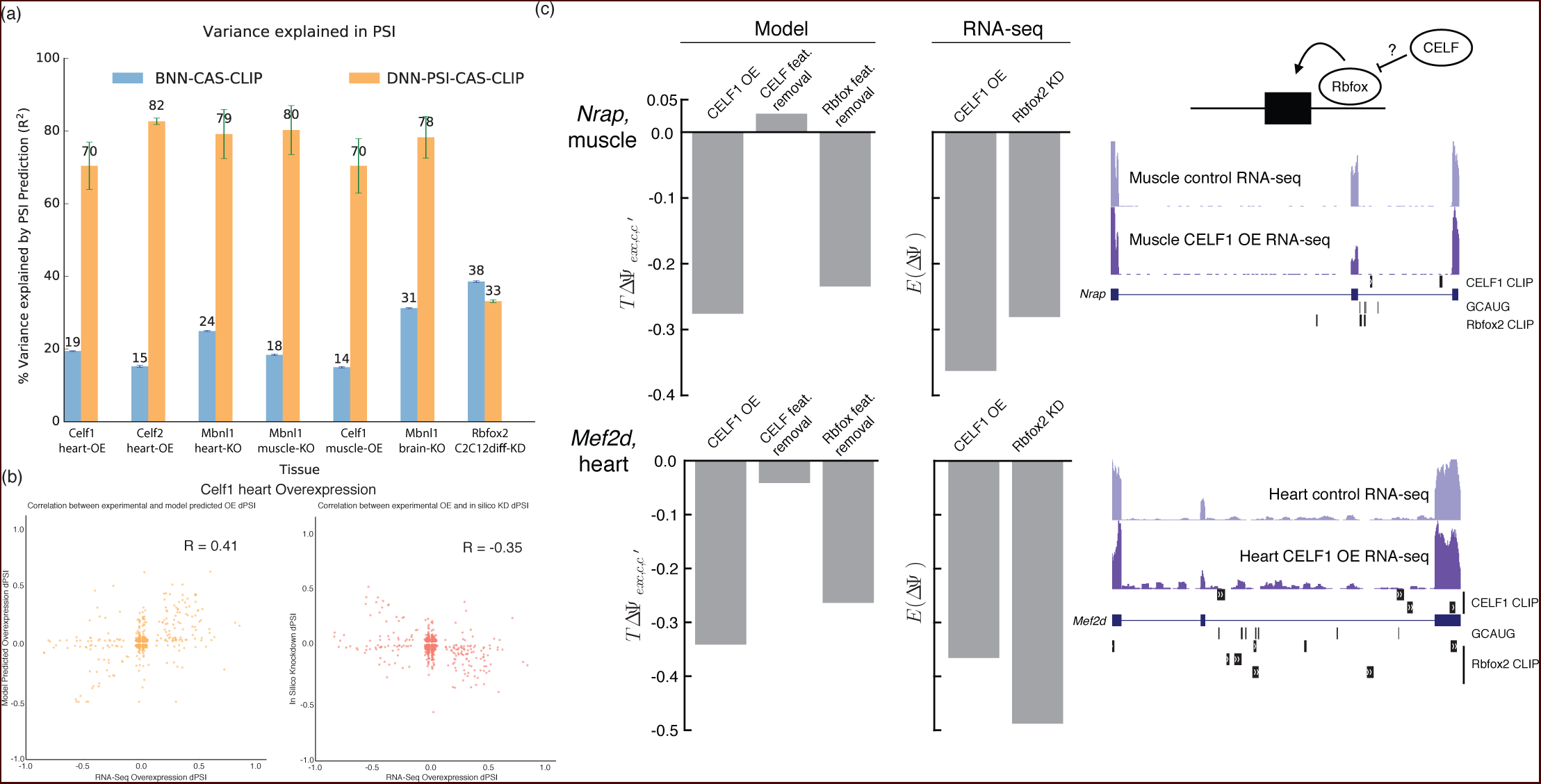
(a) Improvement in % variance explained in PSI for the splice factor modelling for BNN with old target function (blue) versus new model (orange) (b) Correlation plots for Celf1 Overexpression(OE) in mouse heart. Left, showing correlation between experimental and model predicted Overexpression dPSI for Celf1 heart OE. Right, showing correlation between experimental OE and *in silico* KD of Celf. (R: Pearson correlation coe cient) (c) Left: Model predicted changes in exon inclusion for Nrap in muscle (top) or Mef2d in heart (bottom) upon Celf1 overexpression, removal of features related to the Celf family, or removal of features related to the Rbfox family (right bars) as well as quantification of change in inclusion from RNA-Seq upon overexpression Celf1 or knockdown of Rbfox2 in myotubes (left bars). Right: UCSC genome browser view of regulated cassette exons in Nrap (top) and Mef2d (bottom) showing locations of RNA-seq reads in given conditions, Celf1 and Rbfox2 CLIP peaks, and the Rbfox family binding motif GCAUG.

### 3.1 Regulatory Modelling with New Splicing Codes

In order to demonstrate the usefulness of the new splicing codes for splicing regulatory analysis we tested how well the model predicts the effect of splice factor knockdowns on unseen test cases with or without the available KD data. Figure 4(b, left) shows the correlation between the experimental (RNA-Seq) overexpression dPSI and the new model’s predictions in Celf1 heart OE experiment. Good correlation (*R*^2^ = 0.41) indicates that the model learns the effects of overexpression of the splice factor well. Figure 4(b, right) shows the correlation when the model performs *in silico* knockdown of Celf1 by zeroing out the features related to Celf versus the experimental Celf1 overexpression dPSI. Negative correlation (*R*^2^ = −0.35) even without KD data demonstrates how the splicing codes can now accurately predict changes in dPSI with *in silico* knockdowns (For similar plots for the other KO/KD/OE datasets, see Supplementary Figure 1).

Finally, we wished to see if we could gain mechanistic insight into the regulation of physiologically relevant targets in these systems. specifically, exons correctly predicted to have reduced inclusion upon Celf1 over-expression but are not affected by Celf1 related features (Figure 4b,right) are of particular interest in terms of alternative mechanisms of regulation. Two such cases in key genes are shown in Figure 4(c), for the myofibrillar protein *Nrap* [11] in muscle (top) and for the beta microexon in the key myogenic transcription factor *Mef2d* [13] in heart (bottom). quantification using RNA-Seq data from these contexts confirmed the accuracy of the model in predicting Celf1 regulation in both cases (Fig. 4c, compare bars 1 and 4). However, *in silico* removal of Celf related features did not lead to significant changes in exon inclusion in either case(Fig. 4c, compare bars 1 to 2), suggesting indirect regulation could be causing repression upon Celf1 over-expression. In line with this, no Celf1 CLIP peaks were found upstream of these regulated exons (Fig. 4c, right) where Celf proteins have been found to repress exon inclusion [1]. Strikingly, *in silico* removal of features related to the Rbfox family recapitulated the predicted splicing change upon Celf1 overexpression (Fig. 4c, compare bars 1 to 3). Analysis of Rbfox2 knockdown data from myotubes [13] (Fig 4c) or Rbfox1 muscle-specific knockout mice [11] supports that the Rbfox family typically enhances inclusion of these exons. Additionally, a number of Rbfox binding motifs (GCAUG) and CLIP peaks are located just downstream of these exons (Fig 4c, right), where these proteins enhance inclusion [13]. These observations motivated additional study in human T cells where we found Celf2 is a potent repressor of Rbfox2 [6], suggesting that a similar indirect mechanism may be at play in murine muscle and heart where Celf overexpression represses Rbfox proteins to drive splicing changes in these and other targets.

## Discussion

In this study, we offered a new formulation for the task of learning condition specific splicing codes from a compendium of RNA features. First, we introduced a new target function which takes advantage of recent advances in RNA-Seq quantification algorithms [16] and results in a significant accuracy boost for PSI prediction, tissue specific variations, and splice factors target predictions. The new target function allowed us to incorporate samples with missing quantification values or with different degrees of quantification accuracy. This, combined with a pipeline to detect cassette and cassette like exons from RNA-Seq data enabled us to combine many datasets and further improve model accuracy. We also showed how new sources of data for splice factors binding affinity (CLIP-Seq) and regulation (KD/OE experiments) can be integrated to further improve code prediction accuracy.

A known issue with deep models applications for bio-medical studies is their often cryptic nature. However, we were able to demonstrate here how the integrative deep models we developed can be used to gain biological insights for splicing regulation. This included high accuracy of target prediction w/wo available KD/KO experiments, and identifying putative novel regulatory interdependence between splice factors along with the affected targets. We believe this usage demonstration represents only a small portion of the potential of this new breed of models. Future work includes predicting non-cassette splicing variations, robust automated extraction of biological hypotheses from code models, and scaling up to create regulatory codes for many conditions and datasets.

## Acknowledgments

Special thanks to Jorge Vaquero-Garcia for support and advice throughout this project. We would also like to thank NVIDIA Corporation for the kind donation of a Titan X GPU used for this research.

